# Genetic diversity and characterization of circular replication(Rep)-encoding single-stranded (CRESS) DNA viruses

**DOI:** 10.1101/2022.03.09.483724

**Authors:** Perumal Arumugam Desingu, K. Nagarajan

## Abstract

The CRESS-DNA viruses are the ubiquitous virus detected in almost all eukaryotic life trees and play an essential role in the maintaining ecosystem of the globe. Still, their genetic diversity is not fully understood. Here we bring to light the genetic diversity of Replication (Rep) and Capsid (Cap) proteins of CRESS-DNA viruses. We divided the Rep protein of the CRESS-DNA virus into ten clusters using CLANS and phylogenetic analyzes. Also, most of the Rep protein in Rep cluster 1 (R1) and R2 (*Circoviridae, Smacoviridae, Nanoviridae*, and CRESSV1-5) contain the Viral_Rep superfamily and P-loop_NTPase superfamily domains, while the Rep protein of viruses in other clusters has no such characterized functional domain. The *Circoviridae, Nanoviridae*, and CRESSV1-3 viruses contain two domains, such as Viral_Rep and P-loop_NTPase; the CRESSV4 and CRESSV5 viruses have only the Viral_Rep domain, and most of the sequences in the pCRESS-related group have only P-loop_NTPase, and *Smacoviridae* do not have these two domains. Further, we divided the Cap protein of the CRESS-DNA virus into 20 clusters using CLANS and phylogenetic analyzes. The Rep and Cap proteins of *Circoviridae* and *Smacoviridae* are grouped into a specific cluster. Cap protein of CRESS-DNA viruses grouped with one cluster and Rep protein with another cluster. Further, our study reveals that selection pressure plays a significant role in the evolution of CRESS-DNA viruses’ Rep and Cap genes rather than mutational pressure. We hope this study will help determine the genetic diversity of CRESS-DNA viruses as more sequences are discovered in the future.

**Importance:** The genetic diversity of CRESS-DNA viruses is not fully understood. CRESS-DNA viruses are classified as CRESSV1 to CRESSV6 using only Rep protein. This study revealed that the Rep protein of the CRESS-DNA viruses is classified as CRESSV1 to CRESSV6 groups and the new Smacoviridae-related, CRESSV2-related pCRESS-related, Circoviridae-related, and 1 to 4 outgroups, according to the Viral_Rep and P-loop_NTPase domain organization, CLANS, and phylogenetic analysis. Furthermore, for the first time in this study, the Cap protein of CRESS-DNA viruses was classified into 20 distinct clusters by CLANS and phylogenetic analysis. Through this classification, the genetic diversity of CRESS-DNA viruses clarifies the possibility of recombinations in Cap and Rep proteins. Finally, it has been shown that selection pressure plays a significant role in the evolution and genetic diversity of Cap and Rep proteins. This study explains the genetic diversity of CRESS-DNA viruses and hopes that it will help classify future detected viruses.

## Introduction

Circular replication(Rep)-encoding single-stranded (CRESS)-DNA viruses are ubiquitous viruses that are reported to spread worldwide and infect almost all eukaryotic tree of life ^1-3^. CRESS-DNA viruses have also been found in environmental samples such as sewage, seawater, lakes, and springs ^4-11^. Recently, ssDNA viruses have been classified into 13 families ^1^; ten families (*Anelloviridae, Bacilladnaviridae, Bidnaviridae, Circoviridae, Geminiviridae, Genomoviridae, Nanoviridae, Parvoviridae, Redondoviridae*, and *Smacoviridae*) are reported from the eukaryotes ^12^. These viruses are commonly found with replication initiation protein (Rep) and structural capsid protein (Cap) ^1,12^. Of the ten ssDNA virus families found in eukaryotes, the *Bidnaviridae* and *Parvoviridae* families have the linear Genome topology, and the *Anelloviridae* family have a different Rep protein, with the remaining seven families containing circular ssDNA with Rep protein containing the preserved HUH endonuclease motif and superfamily 3 helicase (S3H) domain ^12^.

Recently, these characterized seven families of ssDNA viruses infect eukaryotes (*Bacilladnaviridae, Circoviridae, Geminiviridae, Genomoviridae, Nanoviridae, Redondoviridae*, and *Smacoviridae*), and uncharacterized CRESS-DNA viruses have been classified into separate groups using this characteristic and conserved two-domain Rep protein ^12^. Thus, unclassified CRESS-DNA viruses are classified as CRESSV1 through CRESSV6 ^12^. So far, the Rep protein of CRESS-DNA viruses has been characterized to contain the HUH motif and S3H domain ^1,12^. It is also not widely known what other domains are present in the rep protein of CRESS-DNA viruses that accumulate day by day through metagenomic sequencing in different environmental samples and how they help classify CRESS-DNA viruses. Furthermore, the classification of CRESS DNA viruses by capsid proteins is challenging due to the lack of conserved portions of the capsid proteins of the CRESS DNA viruses as found in the Rep protein ^12^. In particular, the capsid proteins of CRESS DNA viruses are reported to be derived from a number of RNA viruses ^13-16^. It is also largely unknown which of the Cap proteins of the CRESS-DNA viruses that accumulate day by day through metagenomic sequencing in different environmental samples are related to the RNA viruses and the diversity in the Cap proteins of the CRESS-DNA viruses. A recent study found that capsid proteins in Cruciviruses (CRESS DNA virus) are highly conserved and possibly acquired from RNA viruses, but the Rep protein is more diversified than Cap protein ^17^. From these, it is speculated that Cruciviruses may have obtained Rep protein from different CRESS-DNA viruses by recombination ^17^. Therefore, it appears that the genetic variation and recombination of CRESS-DNA viruses can be detected by dividing the capsid proteins of almost identical CRESS-DNA viruses into groups. However, it should be noted that there is no mechanism for classifying the capsid proteins of CRESS-DNA viruses so far.

The present study systematically classified the CRESS-DNA viruses Rep and Cap proteins and reported the presence of different group-specific various domain organizations in the Rep protein. Further, it explains the recombination-mediated evolution of the CRESS-DNA virus and reveals that selection pressure plays a significant role in the evolution of CRESS-DNA viruses’ Rep and Cap genes rather than mutational pressure

## Results

### CLANS based classification of CRESS-DNA Rep protein

As a first step towards understanding the genetic diversity of the CRESS DNA viruses, we analyzed the inter-relationship between the core viral proteins such as Rep and Cap proteins of various isolates of CRESS-DNA viruses. We first chose the Rep protein for our analysis since it shows a high degree of conservation among the CRESS-DNA viruses^2,18-20^. To explore the sequence diversity of the Rep protein of CRESS-DNA viruses, we collected 1160 (sequences details are provided in **Supplementary Data 1)** amino acid sequences of CRESS-DNA viruses from the NCBI Database and grouped them based on pairwise sequence similarity using the CLANS (CLuster ANalysis of Sequences) tool ^21,22^. The analysis grouped the CRESS-DNA virus Rep protein sequences into ten different clusters (R1 to R10) [Rep Cluster 1 (R1)] (a minimum of 10 viral sequences to a maximum of 487 sequences per group) (**Figure 1A**, the individual sequence details in the different clusters are listed in **Supplementary Data 2**). The majority of the clusters, except clusters R7 and R8, showed inter-connections at a *p*-value threshold of 1e^-2^ (**Figure 1A**). Further, we also observed three different superclusters (Super-cluster 1 - clusters R1, R2, R4, and R5; Super-cluster 2 - clusters R3, R6, and R9; Super-cluster 3 - clusters R7 and R8) at a *p*-value threshold of 1e^-5^ (**Supplementary Figure 1A**). For a better understanding of the genetic diversity of the CRESS-DNA virus, we classified the CRESS-DNA viruses into three broad groups as follows (i) culturable CRESS-DNA viruses (*Circoviridae, Geminiviridae, Smacoviridae, Cruciviridae*, etc.)^23^ which are infective, (ii) replication-competent circular DNA (rccDNA) which include the bovine meat and milk factors (BMMF) and Sphinx infective DNA molecule ^24^ and (iii) uncharacterized and uncultivated CRESS-DNA^1^ which were detected as a DNA molecule in the viral metagenomic analysis. Interestingly, most of the sequences in clusters R1 and R2 grouped with highly characterized and culturable viral families of *Circoviridae, Smacoviridae*, and *Cruciviridae*. Further, cluster R8 sequences exclusively belonged to BMMF of rccDNA, while all other clusters included uncultured CRESS-DNA viruses. The remaining clusters (R3, R4, R5, R6, R7, R9, and R10) were classified as uncharacterized and uncultivated CRESS-DNA.

**Figure 1:**
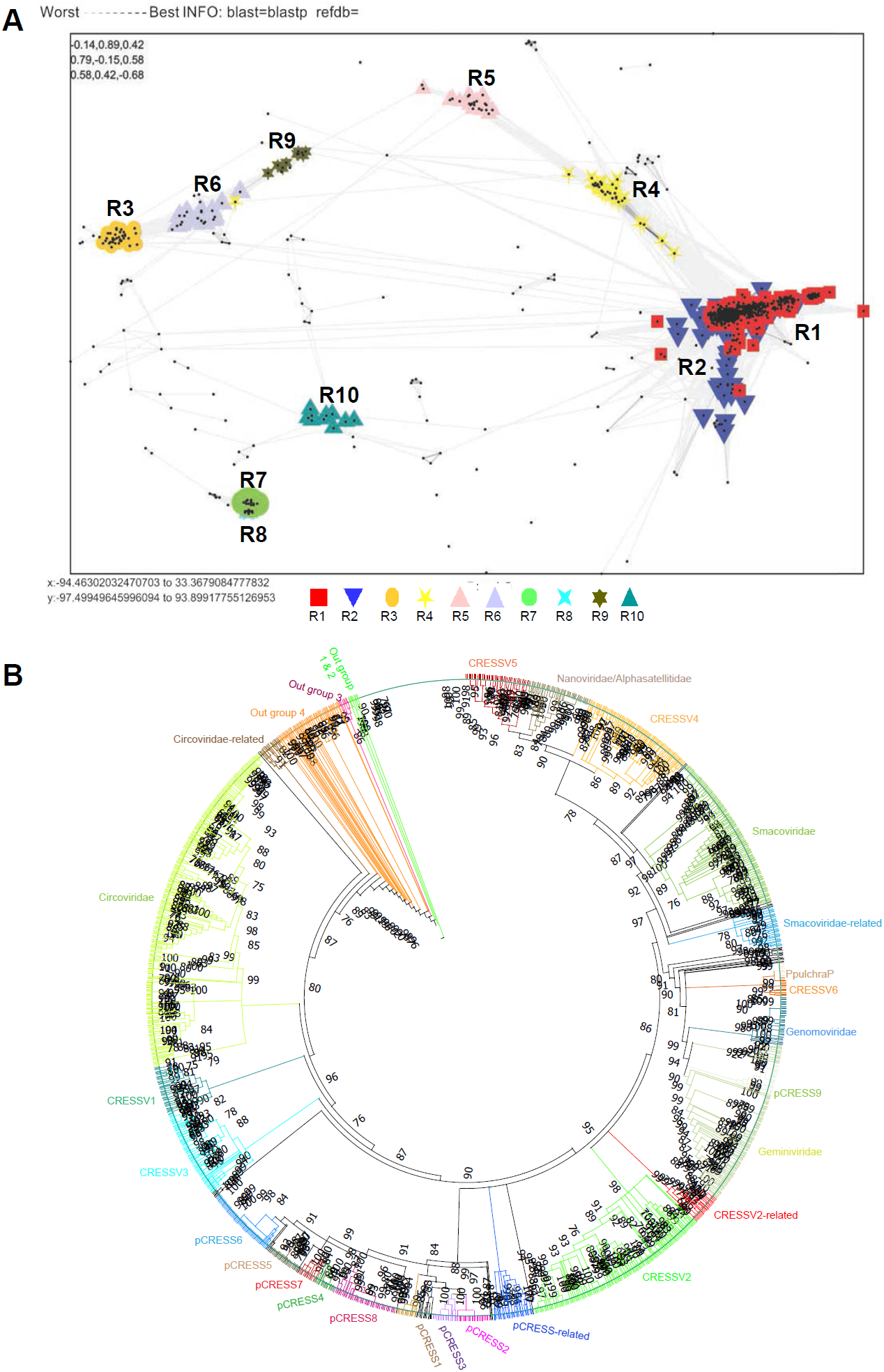
CLANS analysis-based classification of CRESS-DNA virus Rep protein. (**A**) Representative CRESS-DNA virus Rep protein sequences were clustered using CLANS Toolkit by their pairwise sequence similarity network. A total of 1160 amino acid sequences of Rep protein (**Supplementary Data 1**) of CRESS-DNA viruses were used in this analysis Classification of clusters was carried out by a Network-based method using offset values and global average with maximum rounds 10000 in CLANS Toolkit analysis. The *P*-value ≤ 1e^-02^ was used to show the lines connecting the sequences. (**B**) Phylogenetic relationship of Rep protein of CRESS-DNA viruses. The maximum-likelihood method inferred the evolutionary history using the Subtree-Pruning-Regrafting algorithm in PhyML 3.3_1. A total of 1509 amino acid sequences of Rep protein (**Supplementary Data 5**) of CRESS-DNA viruses were used in this analysis.

### Domain organization in the CRESS-DNA virus Rep protein

We were interested to find out if these different Rep protein clusters have any significant differences in the organization of functional domains. The Rep genes of CRESS-DNA viruses have been reported to contain two main functional domains/motif, HUH endonuclease motif and superfamily 3 helicase domains ^1,25^. In this context, we analyzed the domain organization of the Rep protein of viruses from different clusters using the Conserved Domain search tool (https://www.ncbi.nlm.nih.gov/Structure/cdd/wrpsb.cgi?)^26-29^. Interestingly, we found that only the Rep protein of viruses in clusters R1 and R2 displayed functional domains such as Viral_Rep superfamily (Cdd:pfam02407) and P-loop_NTPase superfamily (Cdd:pfam00910). On the other hand, cluster R8 (BMMF) sequences contained Rep_1 superfamily (Cdd:pfam01446), a homologous domain to the Rep1 domain of bacteria involved in plasmid replication. Moreover, we did not find any known putative functional domains in our conserved domain analysis of other clusters consisting of uncultured viruses (Rep cluster R3, R4, R5, R6, R7, R9, and R10). However, it should be noted that R4 and R5 in clusters of Rep protein that do not express these putative functional domains have evolutionary links with R1 and R2 clusters that express functional domains (**Figure 1A**). Similarly, cluster R7 has evolutionary links with cluster R8 that holds the Rep_1 domain (**Figure 1A**).

We then analyzed the diversity of the Rep protein domains in depth belonging to clusters R1 and R2 to further classify the viruses in these clusters, which are highly related to culturable viruses (sequences details are provided in **Supplementary Data 3)**. To explore the different domain organizations present in the viruses of clusters R1 and R2, we re-clustered them into 10 sub-clusters (cluster **a** to **j**) at a *p*-value threshold of 1e^-38^ (**Supplementary Figure 1B**). Of these ten sub-clusters, we noted that sub-clusters such as **a, b, g**, and **h** formed a single group (group 1), and **c, d, i**, and **j** sub-clusters formed a separate group (group 2) (**Supplementary Figure 1B**). Furthermore, it can be seen that there are some evolutionary links between these two groups (group 1 and group 2), but the sub-clusters **e** and **f** together as a separate group (group 3) (**Supplementary Figure 1B**).

The viruses in the sub-cluster **a** majorly contain two main domains: Viral_Rep and P-loop_NTPase domains. Some sequences had one of the following additional domains in between Viral_Rep and P-loop_NTPase domain such as the AAA ATPase domain, Penta-EF hand, DNA-binding ATP-dependent protease La, Type III secretion system protein PrgH-EprH (PrgH), and Parvovirus non-structural protein NS1 (**Supplementary Data 4**). Similarly, cluster **b** viruses also contained the Viral_Rep and P-loop_NTPase domains. In addition, few sequences had a third functional domain between Viral_Rep and P-loop_NTPase domain such as AAA+-type ATPase, SpoVK/Ycf46/Vps4 family, or Type VI protein secretion system component VasK. Moreover, the cluster **b** viruses also contained a combination of domains such as (i) incomplete Viral_Rep + P-loop_NTPase, and (ii) Viral_Rep + incomplete P-loop_NTPase (**Supplementary Data 4**). Also, most sequences in sub-cluster **g** contain Viral_Rep + P-loop_NTPase domains, and some sequences are incomplete with these domains or possess only one of the two domains (**Supplementary Data 4**). Significantly, most sequences in the sub-cluster **h** contain only the P-loop_NTPase domains (**Supplementary Data 4**). More interestingly, it was revealed that most of the sequences in sub-cluster **c** and **d** have only Viral_Rep domains (**Supplementary Data 4**). Also, sub-cluster **i**, which is grouped with sub-cluster **c** and d, contains the Viral_Rep+P-loop_NTPase domains, and sub-cluster **j** contains the Viral_Rep domain+incomplete P-loop_NTPase domains (**Supplementary Data 4**). Finally, it is essential to note that the sequences in sub-clusters **e** and **f** have no known putative functional domains (**Supplementary Data 4**). Collectively, our analyses reveal a vast diversity of domains in the viral Rep protein of CRESS-DNA viruses ranging from lack of any known functional domains to the combination of multiple functional domains.

### Phylogenetic tree based classification of CRESS-DNA virus Rep protein and group-specific domain organization

Recently CRESS-DNA viruses have been classified into different groups CRESSV1 to CRESSV6 using Rep protein ^1,12^. Therefore, we are interested in finding out which CRESSV groups the clusters of Rep protein with varying organizations of domain identified in this current study belong to. To find out, we performed a phylogenetic analysis of the sequences of the Rep protein used in this present study with the sequences used to classify the CRESS-DNA viruses in the previous study ^1^ (**Supplementary Data 5**). In this phylogenetic analysis, we observed that CRESSV6, *P*.*pulchra*, pCRESS9, *Genomoviridae*, and *Geminiviridae* were grouped together, and CRESSV4, CRESSV5, and *Nanoviridae* have formed another group (**Figure 1B**), as in the previous study ^1,12^. As in the previous study ^1^, in plasmid CRESS sequences (pCRESS), CRESS1, pCRESS2, and pCRESS3 formed a separate group, and CRESS4, pCRESS5, CRESS6, pCRESS7, and pCRESS8 formed another group (**Figure 1B**). Furthermore, CRESSV1 and CRESSV3 revealed a close association with *Circoviridae* (**Figure 1B**). In addition, the group that showed a relationship with Smacoviridae was called Smacoviridae-related; the group that showed contact with the CRESSV2 sequences was also called CRESSV2-related; the group that showed a relationship with the pCRESS sequences was called pCRESS-related; the group that showed contact with *Circoviridae* was called Circoviridae-related; also the groups formed an outgroup were named as outgroup 1 to 4 (**Figure 1B**).

We first explored the sub-cluster **a** to **j** created by the clusters R1 and R2 with domain organizations. Notably, we observed the sub-cluster **a** and **b** sequences that revealed the domain organization Viral_Rep+P-loop_NTPase grouped into the CRESSV1, CRESSV2, CRESSV3, *Circoviridae*, and Circoviridae-related groups (**Figure 1B; Supplementary Data 4**). Interestingly, sub-clusters **c** and **d**, which contain only Viral_Rep domains, are grouped with CRESSV4 and CRESSV5, respectively (**Figure 1B; Supplementary Data 4**). We observed that sub-clusters **e** and **f** grouped with *Smacoviridae* without any known putative functional domains (**Figure 1B; Supplementary Data 4**). Significantly, sub-cluster **g**, which display mostly Viral_Rep +P-loop_NTPase domains and some sequences with these domains incomplete or with only one of the two domains, formed the CRESSV2-related group (**Figure 1B; Supplementary Data 4**). Similarly, it should be noted that the sub-cluster **h**, which contains most of the sequences only P-loop_NTPase domains, formed the pCRESS-related group (**Figure 1B; Supplementary Data 4**). Also, sub-cluster **i** often have Viral_Rep+P-loop_NTPase domains and sub-clusters **j** with Viral_Rep domain+incomplete P-loop_NTPase domains grouped with Nanoviridae (**Figure 1B; Supplementary Data 4**).

Next, we explored clusters R3, R4, R5, R6, and R9 without any known putative functional domains. Of these clusters, R4 and R5 combined with clusters R1 and R2 to form Super-cluster 1 (**Supplementary Figure 1A**). Note that cluster R4 forms the Smacoviridae-related group, and cluster R5 forms the outgroup 1 (**Figure 1B; Supplementary Data 4**). We observed that the R3, R6, and R9 clusters formed Super-cluster 2 created outgroup 4, outgroup 3, and outgroup 2, respectively (**Figure 1B; Supplementary Data 4**). These results show that CRESS-DNA virus Rep proteins group into the phylogenetic tree, as is the case with CLANS clustering and domain organizations.

### Classification of CRESS-DNA virus Cap protein using CLANS

While the Rep protein of CRESS-DNA viruses is evolutionarily conserved, the Cap protein is highly diverse ^2,18-20^. Therefore, previous studies analyzed the evolution of capsid proteins primarily by structural fold comparisons rather than sequence comparisons^23,30-32^. However, we took advantage of the recent explosion in the metagenomic data from CRESS-DNA viruses. We employed a sequence comparison method to classify and identify the genetic diversity of CRESS-DNA virus Cap protein. We collected 1823 amino acid sequences of CRESS-DNA viruses from the NCBI Database and grouped them based on pairwise similarity (CLANS analysis) (sequences details are provided in **Supplementary Data 6)**. The analysis classified the CRESS-DNA virus Cap gene sequences into 20 different clusters (minimum of ten sequences per group was considered to classify them as an individual cluster) (the individual sequence details in the different clusters are listed in **Supplementary Data 7**). Most of the clusters show interconnections with other clusters, except the clusters C3 (Cap cluster 3), C4, C19, and C20, which were isolated from other clusters (orphan clusters) in a pairwise similarity network (**Supplementary Figure 2**) (*p*-value threshold of 1e^-02^). Cluster C1 of CRESS-DNA virus sequences clustered with *Circoviridae* viruses, while cluster C2 showed a relationship with *Geminiviridae* viruses, cluster C3 sequences clustered with *Smacoviridae* viruses, and cluster C6 sequences clustered with *Cruciviridae* virus sequences (**Supplementary Data 7**). Among the 20 clusters identified for the Cap protein of the CRESS-DNA viruses (**Figure 2A**), the clusters C2, C7, C8, C11, C12, C15, and C18 form a supercluster (**Figure 2A**) in the sequence similarity network analysis at a *p*-value threshold of >1e^-04^. Similarly, the supercluster consists of clusters C1, C14, and C17 in CLANS analysis (**Figure 2A**; **Supplementary Data 7**). In addition, clusters C3, C4, C5, C6, C9, C10, C13, C16, C19, and C20 were isolated from other clusters (orphan clusters) in a pairwise similarity network (**Figure 2A**; **Supplementary Data 7**) (*p*-value threshold of 1e^- 04^). Collectively, CRESS-DNA virus Cap proteins also split into separate groups in CLANS analysis and are thought to support the classification of Cap proteins.

**Figure 2:**
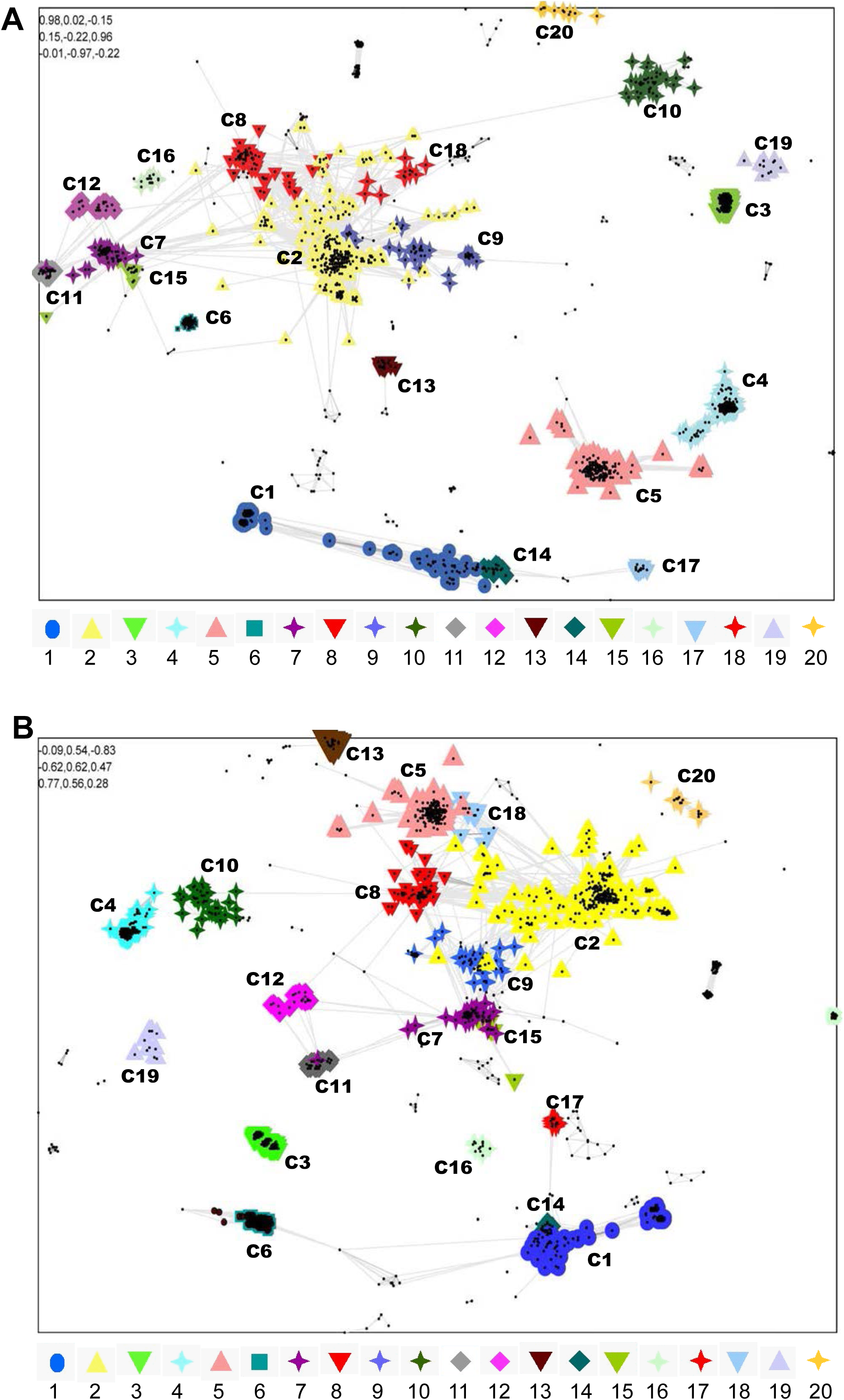
Sequence similarities (CLANS) analysis-based CRESS-DNA virus capsid protein clustering. (**A**) A total of 1823 amino acid sequences of Cap protein of CRESS-DNA viruses (**Supplementary Data 6**) were used and classified by their pairwise sequence similarity network using CLANS. The clusters were classified using the Network-based method using offset values and global average with a maximum of 10000 in CLANS Toolkit analysis. The *P*-value ≤ 1e^−05^ was used to show the lines connecting the sequences. (**B**) Pairwise sequence similarity based on CRESS-DNA virus capsid protein and +RNA viruses relationship. Representative CRESS-DNA virus capsid protein sequences and their relationship with RNA viruses using CLANS Toolkit. A total of 1967 amino acid sequences of Cap protein of CRESS-DNA viruses and +RNA viruses were used in this analysis (**Supplementary Data 8**). The clusters were classified using the Network-based method using offset values and global average with a maximum of 10000 in CLANS Toolkit analysis. The *P*-value ≤ 1e^−02^ was used to show the lines connecting the sequences.

### Only *Cruciviridae* Cap proteins related to RNA viruses

In previous studies, it has been reported that the cap protein of the CRESS-DNA virus is related to the RNA virus ^13-16^, so we were interested to find out which of these 20 clusters is related to the RNA virus. To do this, we retrieved the RNA virus sequences associated with the Cap protein of the CRESS-DNA virus from the NCBI Database and performed CLANS analysis (sequences details are provided in **Supplementary Data 8)**. This analysis noted that RNA viruses revealed association only with *Cruciviridae* virus sequences belonging to cluster C6 at a p-value threshold of 1e-^02^ (**Figure 2B; Supplementary Data 9**)

### Phylogenetic tree based classification of CRESS-DNA virus Cap protein

We examined whether CLANS analysis-based clustering of CRESS-DNA virus cap protein sequences also grouped into the phylogenetic tree. Because cap proteins do not have common domains as seen in CRESS-DNA virus Rep proteins, and the sequence alignments are low from most genetic variants, we performed separate phylogenetic analysis for (i) supercluster C1, C14, C17; (ii) supercluster C2, C7, C8, C11, C12, C15, and C18; and (iii) orphan clusters such as C3, C4, C5, C6, C9, C10, C13, C16, C19, and C20. To do this, we first performed phylogenetic analysis using sequences from the C1, C14, and C17 clusters that formed the Cap protein supercluster. These clusters C1, C14, and C17 are well aligned (**Supplementary Data 10**) and split into separate groups for the phylogenetic tree (**Figure 3A**). Similarly, C2, C7, C8, C11, C12, C15, and C18 clusters are well aligned (**Supplementary Data 11**) and split into separate groups for the phylogenetic tree (**Figure 3B**). In particular, C8, C11, and C12 formed an outgroup, and this outgroup group C8 was somewhat detached, and C11 and C12 grouped slightly closer together into the phylogenetic tree (**Figure 3B**) as seen in the CLANS analysis (**Figure 2A**). Similarly, the C7 and C15 clusters grouped in the phylogenetic tree (**Figure 3B**), as seen in the CLANS analysis (**Figure 2A**), and the C18 and some C2 sequences grouped together with this (C7 and C15) group (**Figure 3B**). Also, although the cluster C2 sequences are majorly grouped together, it is noteworthy that some sequences are grouped together with a group formed by C8, C11, and C12 and a group created by C7, C15, and C18 (**Figure 3B**). We then performed phylogenetic analysis separately for the orphan clusters C3, C4, C5, C6, C9, C10, C13, C16, C19, and C20 clusters. Thus, the sequences in these clusters are well-aligned C3 (**Supplementary Data 12**), C4 (**Supplementary Data 13**), C5 (**Supplementary Data 14**), C6 (**Supplementary Data 15**), C9 (**Supplementary Data 16**), C10 (**Supplementary Data 17**), C13, C16, C19 and C20 (**Supplementary Data 18**), to form the phylogenetic tree C3 (**Supplementary Figure 3A**), C4 (**Supplementary Figure 3B**), C5 (**Supplementary Figure 4A**), C6 (**Supplementary Figure 4B**), C9 (**Supplementary Figure 5A**), C10 (**Supplementary Figure 4B**), C13, C16, C19, and C20 (**Supplementary Figure 5C**).

**Figure 3:**
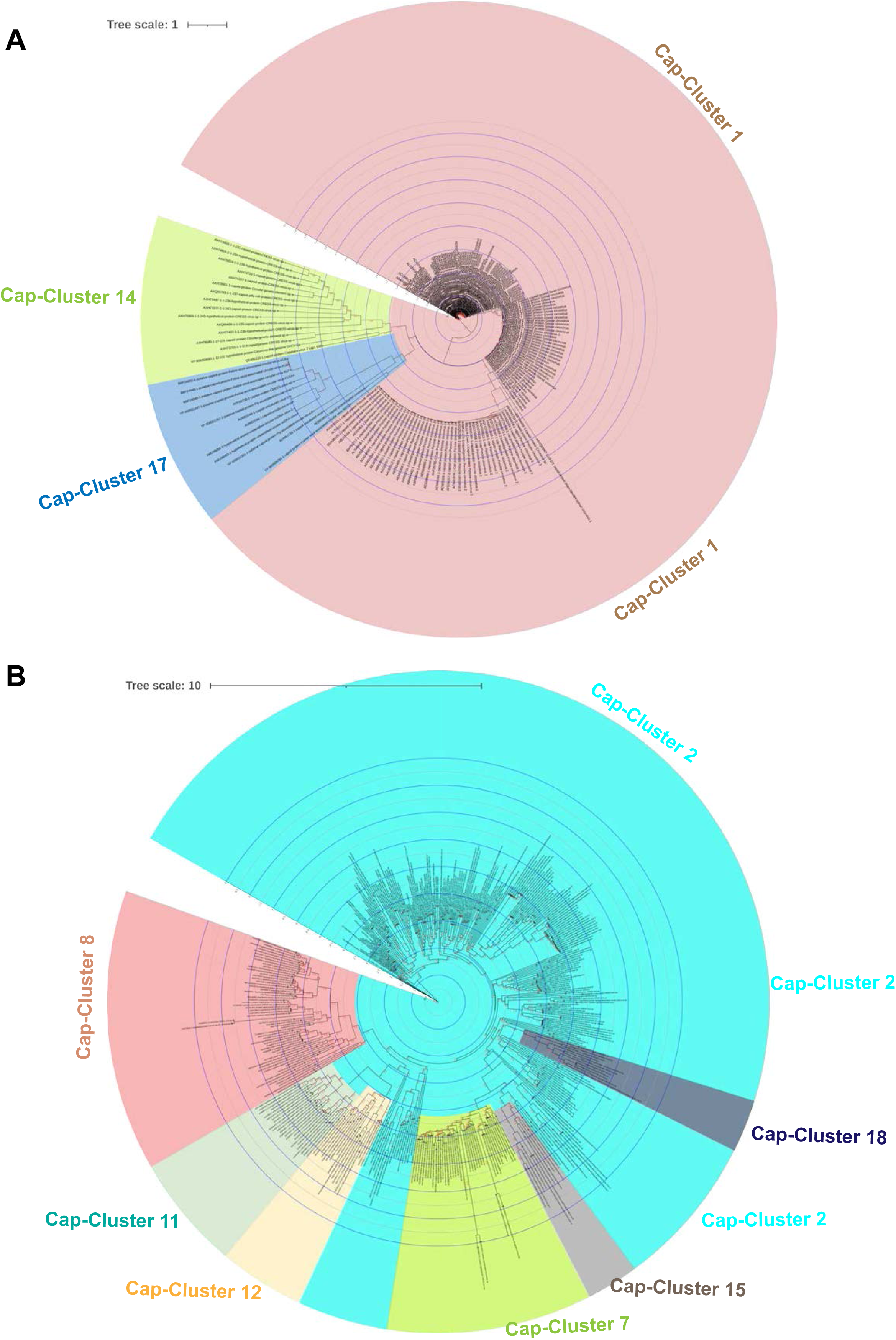
Phylogenetic relationship of CRESS-DNA virus Cap protein superclusters. (**A**) Phylogenetic tree depicting the genetic relationship between the CRESS-DNA virus Cap protein supercluster formed by the clusters C1, C14, and C17. The details of sequences in each cluster (**Supplementary Data 7**) and alignment are provided in **Supplementary Data 10**. (**B**) The phylogenetic tree represents the genetic relationship between the CRESS-DNA virus Cap protein supercluster created by the clusters C2, C7, C8, C11, C12, C15, and C18. The details of sequences in each cluster (**Supplementary Data 7**) and alignment are provided in **Supplementary Data 11**. The maximum-likelihood method inferred the evolutionary history using the Subtree-Pruning-Regrafting algorithm and bootstrap values in PhyML 3.3_1.

### Recombination mediated evolution of CRESS-DNA viruses

Recently, it has been reported that the cap protein of Cruciviruses is very similar, but the Rep protein may be derived from different sources with greater diversity ^17^; we examined whether the sequences in the cluster of these 20 Cap proteins received the Rep protein from the same group or from different groups. To do this, we took the representative sequences in each Cap-cluster and identified the phylogenetic tree groups that contain its Rep protein (**Supplementary Data 19**). In this analysis, it appears that the sequences in the same cap-cluster have different groups of rep proteins (**Supplementary Data 19**). From these, it can be inferred that the CRESS-DNA virus has the potential to acquire genetic diversity through recombination in the Cap and Rep genes.

### Role of host codon usage selection pressure on Rep gene evolution

Since we observe homology at amino acid levels between the Rep gene of CRESS-DNA viruses but not any significant identity at the nucleotide sequence level, we suspected this might be due to this virus’s host codon usage bias-based selection pressure. To explore this, we first analyzed the base composition of 1115 nucleotide sequence of CRESS-DNA viruses’ Rep genes (the details of nucleotide sequences used in the analysis are presented in **Supplementary data 20**) as AT to GC ratio can affect codon usage in microbes ^33,34^. Our study revealed that the Rep gene of CRESS-DNA viruses contains A>T>G>C with AT%>GC% (average GC content is 43.6±SD7.06) (**Figure 4A; Supplementary Data 21**). We next analyzed the codon usage bias using the effective number of codon usage (ENc) analysis. ENc values <35 indicate high codon bias, and values >50 show general random codon usage^35,36^. The Rep gene of CRESS-DNA viruses has ENc values ranging from 31 to 61, while most of the ENc values fall between 40 and 60 (average ENc 51.004±SD5.73) (**Figure 4B; Supplementary data 21**), indicating weak to strong codon usage bias. In addition, we calculated the relative synonymous codon usage (RSCU) value which is the ratio between the observed to the expected value of synonymous codons for a given amino acid. A RSCU value of one indicates that there is no bias for that codon. In contrast, RSCU values >1.0 have positive codon usage bias (defined as abundant codons), and RSCU values <1.0 have negative codon usage bias (defined as less-abundant codons) ^36,37^. Our analysis revealed that the RSCU values of 28 codons were >1 and 31 codons were <1 in the Rep gene of all the CRESS-DNA viruses (**Figure 4C; Supplementary Data 21**), clearly indicating a codon usage bias (both positive and negative).

**Figure 4:**
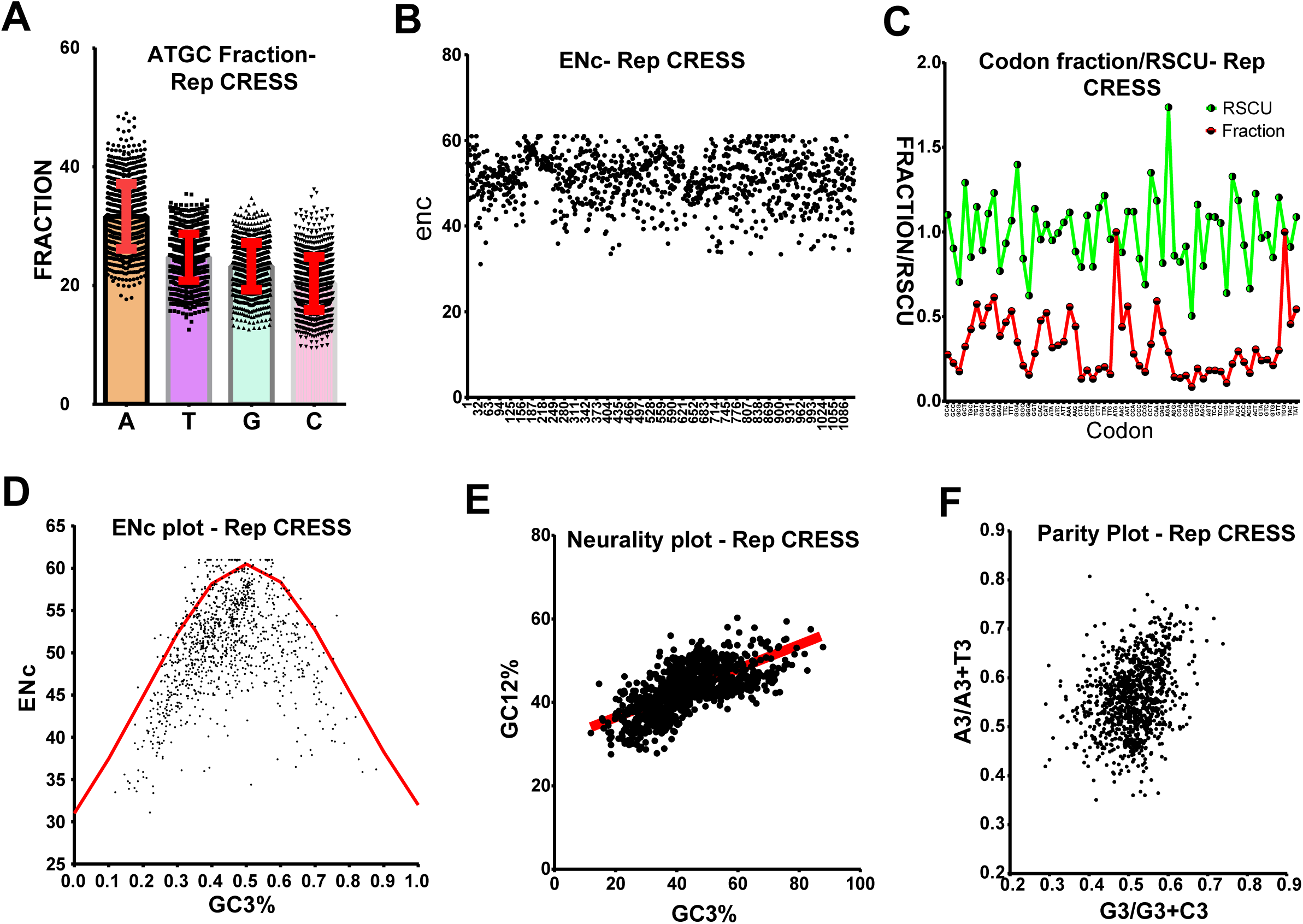
Host codon usage selection pressure on Rep gene of CRESS-DNA virus evolution. (**A**) Representing the A, T, G, and C fraction; (**B**) Represent the ENc values; (**C**) represent the codon usage fraction and RSCU values; (**D**) Represents ENc plotted against GC3s; (**E**) Neutrality plot analysis of the GC12 and that of the GC3; and (**F**) Parity Rule 2 (PR2)-bias plot (Total of 1115 nucleotide sequences of Rep gene of CRESS-DNA viruses were used in this analysis).

Next, we performed ENc-GC3s plot analysis where the ENc values are plotted against the GC3s values (GC content at the third position in the codon) to determine the significant factors such as selection or mutation pressure affecting the codon usage bias^38^. In this analysis, genes whose codon bias is affected by mutations will lie on or around the expected curve. In contrast, genes whose codon bias is affected by selection and other factors will lie beneath the expected curve ^36,38^. Interestingly, we observed that most of the points fall below the expected curve in the ENc-GC3s plot analysis (**Figure 4D; Supplementary Data 21**), indicating the strong presence of selection pressure rather than mutation pressure. Similarly, neutrality plot analysis where GC12 values (average of the GC content percentage at the first and second position in the codon) are plotted against GC3 values to evaluate the degree of influence of mutation pressure and natural selection on the codon usage patterns, displayed a slope of 0.2899 (Y=0.2899*X+30.61, r= 0.662; p<0.0001) (**Figure 4E; Supplementary Data 21**), indicating that the mutation pressure and natural selection were 28.9% and 71.1%, respectively. Moreover, we performed Parity rule 2 bias analysis, where the AT bias [A3/(A3+ T3)] is plotted against GC-bias [G3/(G3 + C3)] to determine whether mutation pressure and natural selection affect the codon usage bias^38^. If A = T and G = C, it indicates no mutation pressure and natural selection, while any discrepancies indicate mutation pressure and natural selection. Our analysis of CRESS-DNA Rep gene sequences shows unequal A to T and G to C numbers, indicating the presence of mutation and selection pressure (**Figure 4F, Supplementary Data 21**). Taken together, these results suggest that CRESS-DNA has wide host-range adaptation, maintaining better codon usage pattern with bacteria, and further selection pressure has played a significant role in the evolution of the CRESS-DNA viruses Rep gene rather than mutational pressure.

### Role of host codon usage selection pressure on Cap gene evolution

Similar to the Rep gene, our NCBI nucleotide BLAST analysis of the Cap gene also showed limited homology between the Cap gene of CRESS-DNA viruses. Since we observed a strong codon-bias-based evolution in the Rep gene of CRESS-DNA viruses (**Figure 4A-F**), we tested whether the Cap gene of the CRESS-DNA viruses also shows codon-bias-based evolution to explore whether the evolution of Cap gene was influenced by mutation pressure or selection pressure, we retrieved 1134 nucleotide sequences of Cap genes of CRESS-DNA viruses (**Supplementary Data 22**) from NCBI public database. Our analysis of the nucleotide base composition of the Cap gene revealed that the Cap gene contains AT%>GC% (average GC content is 44.72±SD 5.96) (**Figure 5A; Supplementary Data 23**). Further, the Cap protein of the CRESS-DNA virus has ENc value ranging from 33 to 61, while most of the sequence ENc values fall between 40 to 60 (average ENc 51.54±SD 5.36) (**Figure 5B; Supplementary Data 23**). Similarly, the RSCU values of 27 codons were >1, and 31 codons were <1 in all the Cap genes of CRESS-DNA viruses (**Figure 5C; Supplementary Data 23**). Also, nine codons showed RSCU values <0.7, and 6 codons showed RSCU values>1.5, indicating the presence of under-represented and over-represented codon bias in the Cap gene, respectively (**Supplementary Data 23**). Moreover, we performed ENc-GC3s plot analysis and found that most points fall below the expected curve in the ENC-GC3s plot (**Figure 5D; Supplementary Data 23**). In line with this, the neutrality plot displayed a slope of 0.1404 (Y=0.1404*X+40.64; r= 0.512; p<0.0001) (**Figure 5E; Supplementary Data 23**), indicating 14% of mutation pressure and 86% of selection pressure in this gene and Parity rule 2 bias analysis showed discrepancies in the A to T and G to C numbers in the third position of the codon (**Figure 5F; Supplementary Data 23**). Taken together, these results indicate that the selection pressure played a more significant role in the Cap gene than the Rep genes of CRESS-DNA viruses.

**Figure 5:**
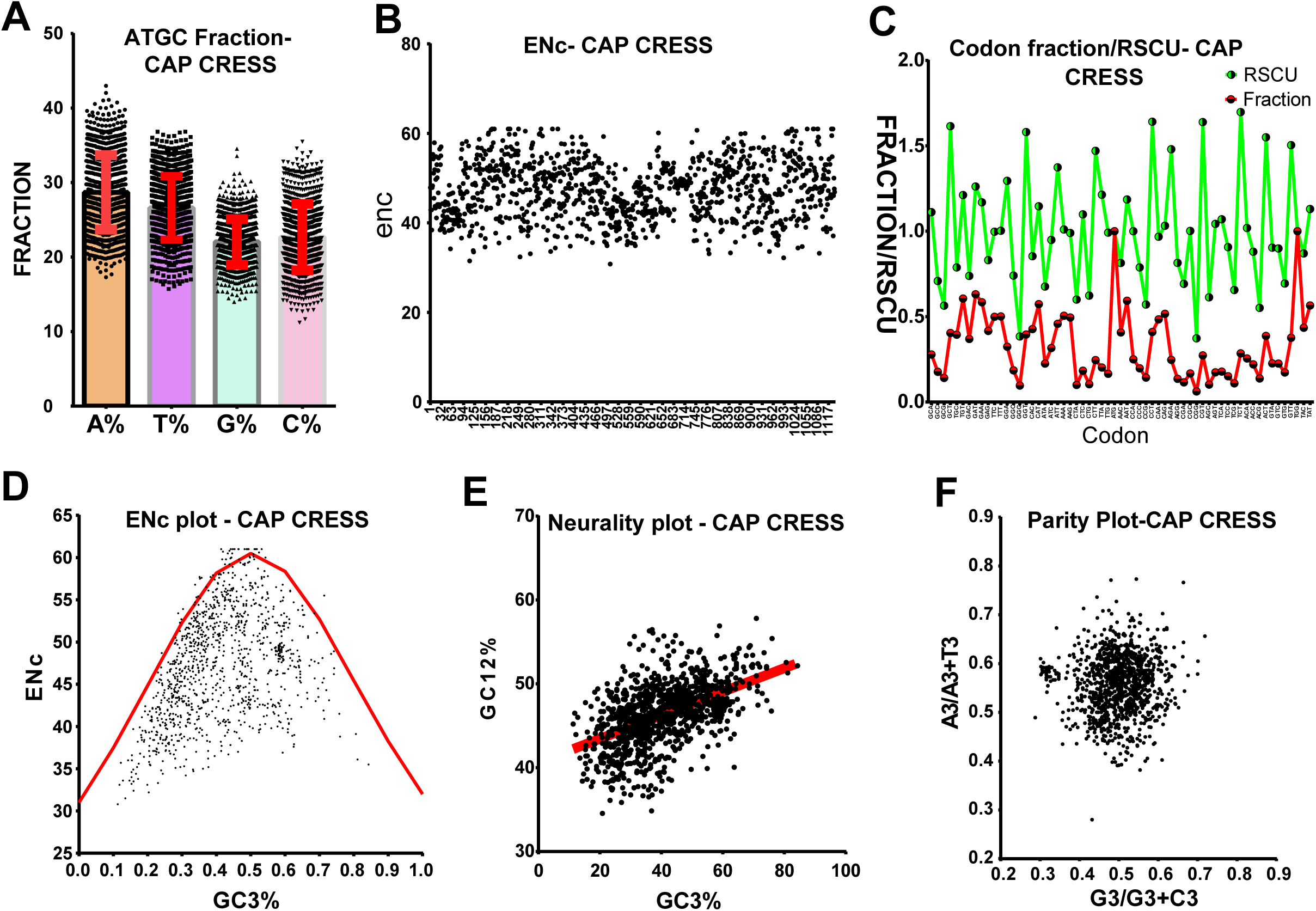
Host codon usage selection pressure on Cap gene of CRESS-DNA virus evolution. (**A**) A, T, G, and C fraction; (**B**) ENc values; (**C**) Codon usage fraction and RSCU values; (**D**) ENc plotted against GC3s; (**E**) Neutrality plot analysis of the GC12 and that of the GC3; and (**F**) Parity Rule 2 (PR2)-bias plot (Total of 1134 nucleotide sequences of Cap gene of CRESS-DNA viruses used in this analysis).

## Discussion

The genetic diversity of CRESS-DNA viruses so far is known only to be the tip of the iceberg. Many novel CRESS-DNA viruses have recently been detected by metagenomic sequencing ^8,39-41^. The rapid development of metagenomic sequencing suggests that in the future, most CRESS-DNA viruses will be detected from different sources and that these CRESS-DNA viruses will be divided into different virus families. Therefore, it is hoped that identifying and classifying genetic diversity in CRESS-DNA viruses will help determine their importance in transmission and pathogenesis and design antivirals and vaccines for appropriate control and prevention. However, the classification of CRESS-DNA viruses has been determined using only the Rep protein ^1,12^. This is because Rep protein contains conserved HUH motif, and S3H domains, while Cap protein is unclassified because it has high genetic diversity without being conserved ^1,12^. However, of the cruciviruses that classify Cap protein well, the report that Cap proteins are nearly identical and that the highly diverse Rep protein may be derived from different CRESS-DNA virus sources is critical here ^17^. Therefore, it can be expected that the genetic diversity and genetic recombination events of CRESS-DNA viruses can be determined by detecting and classifying the diversity in both Rep and Cap proteins.

It is noteworthy that recently, unclassified CRESS-DNA viruses using the Rep protein of CRESS-DNA viruses were grouped into six groups called CRESSV1 to CRESSV6 ^1,12^. The present study reveals that there are not only CRESSV1 to CRESSV6 groups but also groups with Smacoviridae-related, CRESSV2-related, pCRESS-related, Circoviridae-related, and 1 to 4 outgroups are there. So far, it has been reported that the Rep protein of the CRESS-DNA virus contains the HUH motif and S3H domains ^1,12^. In this study, we report the presence of domains such as Viral_Rep superfamily (Cdd: pfam02407) and P-loop_NTPase superfamily (Cdd: pfam00910) in the Rep protein of most CRESS-DNA viruses. Furthermore, this present study revealed the presence of these two domains in the CRESSV1, CRESSV2, CRESSV3, *Circoviridae*, and Circoviridae-related groups and the *Nanoviridae* group. However, CLANS and phylogenetic analyses clarify the viral_Rep and P-loop_NTPase domains in the CRESSV1, CRESSV2, CRESSV3, Circoviridae, and Circoviridae-related groups are very close and distinct from the *Nanoviridae* group. It is noteworthy that CRESSV1, CRESSV2, CRESSV3, Circoviridae, and Circoviridae-related groups together formed the Rep sub-cluster **a** and **b** and the sequences in the *Nanoviridae* group Rep sub-cluster **i** and **j** (**Supplementary Figure 1B**). Our phylogenetic tree (**Figure 1B**) and previous study ^12^ reflect this diversity. In particular, some sequences in the rep sub-cluster **a** and **b** appear to have an additional domain (Penta-EF hand, DNA-binding ATP-dependent protease La, Type III secretion system protein PrgH-EprH (PrgH), etc.) between the Viral_Rep and P-loop_NTPase domains. From the acquisition of such additional functional domains, it is clear that these viruses are stepping into the next stage of evolution, and when more sequences are found later, it is possible to speculate that they are likely to be classified as separate virus families. However, since these sequences are detected by metagenomic sequencing from uncultured viruses and maybe sequence alignment error, it may be imperative to isolate the viruses and identify the significance of these additional functional domains.

Furthermore, in CLANS analysis, the sub-cluster **c** and **d** were grouped with the sub-cluster **i** and **j** reacting with *Nanoviridae* (Suppl**ementary Figure 2B**), of which the sub-cluster **c** was CRESSV4, and the sub-cluster **d** were CRESSV5 viruses (**Supplementary Data 4**); and reflect in our phylogenetic tree (**Figure 1B**) and previous study ^12^. In particular, the CRESSV4 and CRESSV5 viruses have only the Viral_Rep domain, and the sequences in the sub-cluster **j** related to Nanoviridae are the Viral_Rep+incomplete P-loop_NTPase, and the sequences in the sub-cluster **i** are the Viral_Rep+P-loop_NTPase domains. Of these, it can speculate that the CRESSV4 and CRESSV5 viruses, which have only the Viral_Rep domain only, may have appeared first, followed by the sub-cluster **j** with the viral_Rep+incomplete P-loop_NTPase, and finally the sub-cluster **i** virus with the Viral_Rep+P-loop_NTPase domains. Similarly, sub-cluster **g** (CRESSV2-related group) that are often Viral_Rep+P-loop_NTPase domains and some sequences where these domains are incomplete or show only one of the two domains may have led to the emergence of CRESSV2 viruses with Viral_Rep+P-loop_NTPase domains. Furthermore, it is essential to note that sub-cluster **h**, which usually contains only the P-loop_NTPase domain, formed the pCRESS-related group. Interestingly, no functional domains were found in Smacoviridae’s Rep protein in the Conserved Domain search tool, which revealed links between *Circoviridae* and *Nanoviridae* in CLANS and biogenetic analyzes. Similarly, no functional domains were found in the sequences in group R5 (CLANS) or Smacoviridae-related group (phylogenetic tree). It is noteworthy that the sequences of Rep protein that formed the outgroups in this phylogenetic analysis are the clusters of CLANS analysis, R3, R5, R6, and R9, forming separate groups. R3, R5, R6, and R9 clusters formed Super-cluster 2 created outgroup 4, outgroup 1, outgroup 3, and outgroup 2, respectively. Remarkably, no functional domains are found in the sequences in the R3, R5, R6, and R9 clusters that make up the outgroups. However, the sequences that make up the outgroups are detected from uncultured viruses by metagenomic sequencing, and it can be expected that the functional significance will be revealed by isolating these viruses and characterizing the Rep protein.

Cap protein was high in genetic diversity, making it challenging to align and phylogenetically classify correctly. Therefore, in this study, we subdivided the closest sequences into clusters using CLANS analysis and then did phylogenetic classification by aligning them well using the corresponding clusters. First, in this study, the Rep proteins were divided into clusters in the CLANS analysis and then phylogenetic classification using the related clusters, which is consistent with phylogenetic classification in the previous studies ^1,12^. Accordingly, we divide the cap protein into 20 clusters using CLANS analysis, and (i) supercluster C1, C14, C17; (ii) supercluster C2, C7, C8, C11, C12, C15, and C18; and (iii) orphan clusters such as C3, C4, C5, C6, C9, C10, C13, C16, C19, and C20 became well aligned and led to phylogenetic classification. Furthermore, only the Cruciviridae virus sequences in Cap-cluster C6 revealed evolutionary relationships with RNA viruses, but future studies need to determine the evolutionary origins of the sequences in other Cap-clusters. Remarkably, this study revealed that viruses in the same Cap-cluster derive their Rep protein from groups of different Rep proteins, which can be speculated to be generated by genetic recombination. These can be believed to underscore the importance of classifying Cap protein. Finally, this study makes it clear that selection pressure plays a more significant role than mutational pressure in the genetic diversity and evolution of CRESS-DNA virus Cap and Rep protein. Therefore, it can be expected that there will be more opportunities to detect CRESS-DNA viruses with greater genetic diversity and/or recombination in the future. We hope this study will help determine the genetic diversity/recombination of CRESS-DNA viruses as more sequences are discovered in the future.

In conclusion, to the best of our knowledge, this is the first report on the CRESS-DNA virus Rep protein classification using a different domain organization pattern; and CLANS and phylogenetic analysis based on the classification of Cap protein. Furthermore, this study also clarifies the genetic diversity in CRESS-DNA viruses formed by recombination and selection pressures in Cap and Rep proteins. It is widely expected that CRESS-DNA viruses, which have tremendous genetic diversity in the future, will be able to be detected from different sources in different parts of the world through rapidly growing metagenomic sequences. We hope this study will help you determine and accurately classify using CLANS, phylogenetic groups, the domain organization pattern, genetic diversity, and recombination of those CRESS-DNA viruses.

## Materials and Methods

### I. Databases search, collection, and curation

Complete genome sequences of CRESS-DNA viruses were retrieved from the NCBI nucleotide database (https://www.ncbi.nlm.nih.gov/nucleotide/). Rep and Cap genes’ characterized protein-coding sequence (CDS) region and their corresponding amino acid sequences were retrieved from the database in the available complete genome sequence of CRESS-DNA viruses. The uncharacterized CDS of CRESS-DNA viruses were classified as a Cap/Rep protein using NCBI protein BLAST (https://blast.ncbi.nlm.nih.gov/Blast.cgi?PAGE=Protein) analysis, and the sequences were retrieved. Further, complete genome sequences which contain Cap/Rep of every CRESS-DNA were individually used to perform separate NCBI protein BLAST analysis (https://blast.ncbi.nlm.nih.gov/Blast.cgi?PAGE=Protein). Their BLAST aligned sequences of other ssDNA viruses (example: Circoviridae, Smacoviridae, Cruciviridae, etc.) were retrieved.

### II. CLANS (CLuster ANalysis of Sequences) analysis

The CLANS analysis was performed in the online Toolkit software (https://toolkit.tuebingen.mpg.de/tools/clans). The protein sequences retrieved from the NCBI database were subjected to the pairwise sequence similarity calculation using the online CLANS analysis in the Toolkit^21^ with a scoring matrix of BLOSUM45 and BLAST HSP’s (High Scoring Pair) up to an E-value of 1e^-2^. Next, the CLANS files obtained from the Toolkit were visualized in a Java application (clans.jar)^22^. A minimum of 1,00,000 rounds was used to show the sequences connection and clusters in the clans.jar application. The clusters were classified based on the Network method using offset values and global average with maximum rounds of 10000 in clans.jar analysis.

### III. Analysis of functional domain organization in the protein

We determined the domain organizations in the Rep protein of the CRESS-DNA virus using the Conserved Domain search tool (https://www.ncbi.nlm.nih.gov/Structure/cdd/wrpsb.cgi). For this, we used the CDD v3.19-58235 PSSms database, Expect Value threshold line 0.01, Composition-based statistics adjustment applied, and Performed by the maximum number of hits to 500 in the Conserved Domain Search tool ^26-29^.

### IV. Phylogenetic analyses

The phylogenetic analysis was performed in PhyML 3.3_1 using the amino acid sequences of the Rep/Cap protein of the CRESS-DNA virus clustered into clusters in the CLANS analysis, which was retrieved from the NCBI public database. The phylogenetic analysis is in PhyML 3.3_1, Evolutionary model LG, Equilibrium frequencies Empirical ML-Model, discrete gamma model [number of categories (n=4)], tree topology search with SPR (Subtree Pruning and Regraphing), tree topology, branch length, and model parameters are optimizing parameters, and SH-like statistics are used to test the branch support ^42-44^. Further, the phylogenetic trees were visualized through the interactive tree of life (iTOL) v5 ^45^.

### V. Codon usage bias analysis

#### a) Nucleotide sequence composition analysis

The nucleotide composition of CDSs, specifically the A%, T%, G%, and C% composition of the Rep/Cap genes of CRESS-DNA viruses, were analyzed using Automated Codon Usage Analysis (ACUA) Software^46^.

#### b) Relative Synonymous Codon Usage (RSCU) Analysis

RSCU value is the ratio between the observed to the expected value of synonymous codons for a given amino acid. When the RSCU value is one, it indicates that there is no bias for that codon^36,37^. This study determined the RSCU values using the ACUA Software^46^. The nucleotide sequences of Rep/Cap genes of CRESS-DNA viruses obtained from the NCBI nucleotide public database were used for this analysis.

#### c) Effective Number of Codons (ENc)

The effective number of codon usage from 61 codons for the 20 amino acids is one method that determines the codon usage bias and may range from 20 to 61. ENc values <35 indicate high codon bias, and values >50 show general random codon usage^35,36^. In this study, the ENc values were determined on the online server (http://ppuigbo.me/programs/CAIcal/)^47^, and the input nucleotide sequences used in the CAI calculation were used in this analysis.

### VI. Determining the selection and mutation pressure

#### a) ENc-GC3s plot

In this analysis, the ENc values are plotted against the third position of GC3s of codon values to determine the significant factors such as selection or mutation pressure affecting the codon usage bias^38^. The expected curve was determined by estimating the expected ENc values for each GC3s as recommended in previous publications^36,38^. The ENc and GC3s for every gene were obtained from an online CAI analysis server (http://ppuigbo.me/programs/CAIcal/)^47^. The genes would lie on or around the expected curve when mutation pressure only affects codon bias. In contrast, they would fall considerably below the expected curve if codon bias is influenced by selection and other factors^36,38^.

#### b) Neutrality plot analysis

In a neutrality plot, GC12 values of the codon are plotted against GC3 values to evaluate the degree of influence of mutation pressure and natural selection on the codon usage patterns. The GC12 and GC3 values for the nucleotide sequences of Rep/Cap genes of CRESS-DNA viruses were obtained from an online CAI analysis server (http://ppuigbo.me/programs/CAIcal/)^47^.

#### c) Parity Rule 2 (PR2)-bias plot

The PR2-bias, the AT bias [A3/(A3+ T3)] is plotted against GC-bias [G3/(G3 + C3)] to mutation pressure and natural selection affecting the codon usage bias^38^. The A3, T3, G3, and C3 values of nucleotide sequences of Rep/Cap genes of CRESS-DNA viruses were obtained using the ACUA Software^46^.

## Acknowledgments

PAD is a DST-INSPIRE faculty is supported by research funding from the Department of Science and Technology (DST/INSPIRE/04/2016/001067), Government of India, and Core grant from the Science and Engineering Research Board (SERB) (CRG/2018/002192), Department of Science and Technology (DST), Government of India.

## Data Availability Statement

We have retrieved the nucleotide sequences from publically available NCBI databases. Further, all the nucleotide sequences accession numbers and names are indicated in the respective figures and supplementary data.

## FIGURE LEGENDS

**Supplementary Figure 1:**
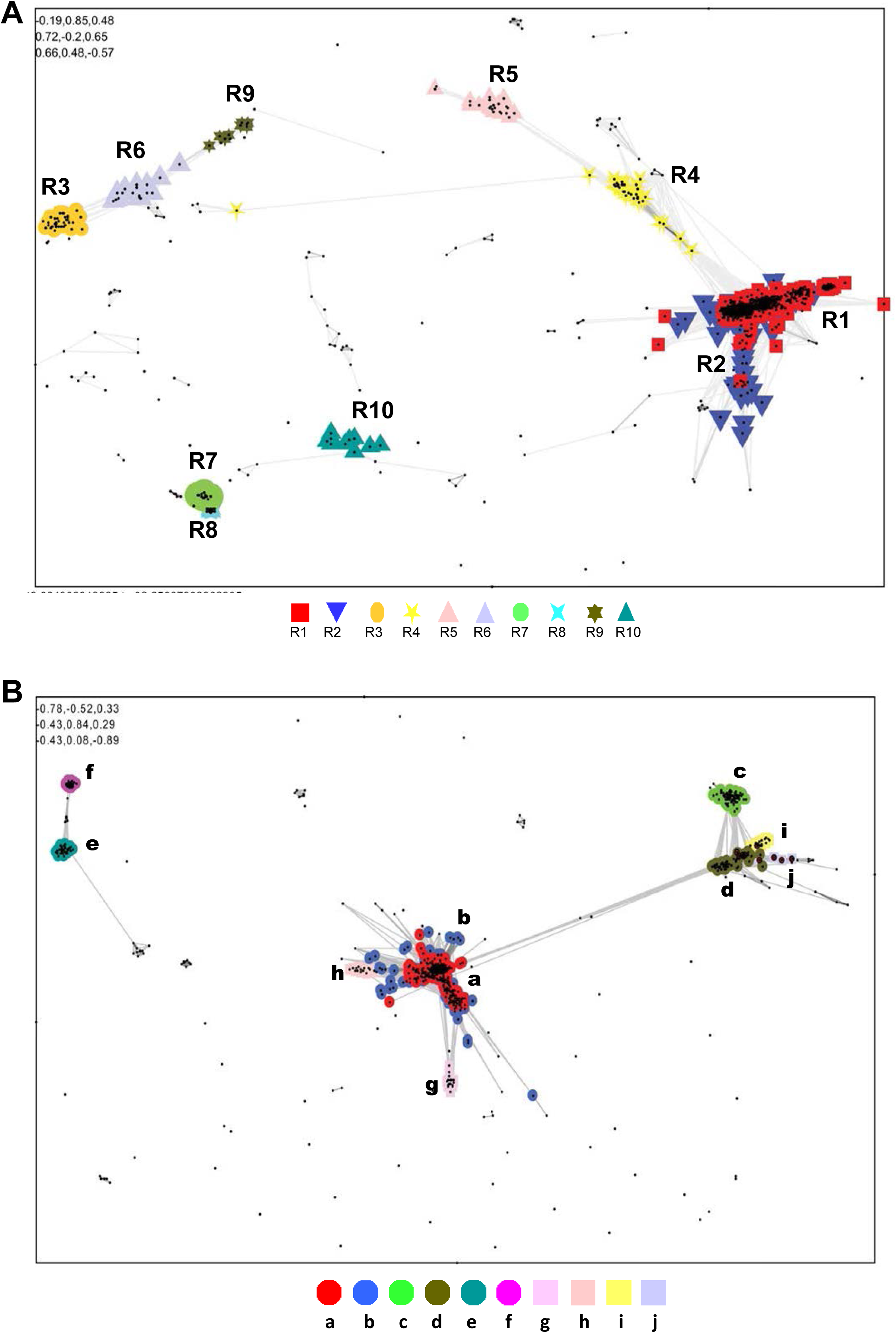
Pairwise amino acid sequence similarity network-based CRESS-DNA virus classification of Rep protein of CRESS-DNA viruses. (**A**) A total of 1160 amino acid sequences of Rep protein of CRESS-DNA viruses (**Supplementary Data 1**) were clustered by CLANS. The clusters were classified using the Network-based method using offset values and global average with a maximum of 10000 in CLANS Toolkit analysis. The *P*-value ≤ 1e^-05^ was used to show the lines connecting the sequences. (**B**) Sub-clustering of the cluster 1 and 2 protein sequences of CRESS-DNA viruses Rep proteins into 10 different clusters (cluster **a** to **j**) at a *P*-value threshold of 1e^-38^.

**Supplementary Figure 2:**
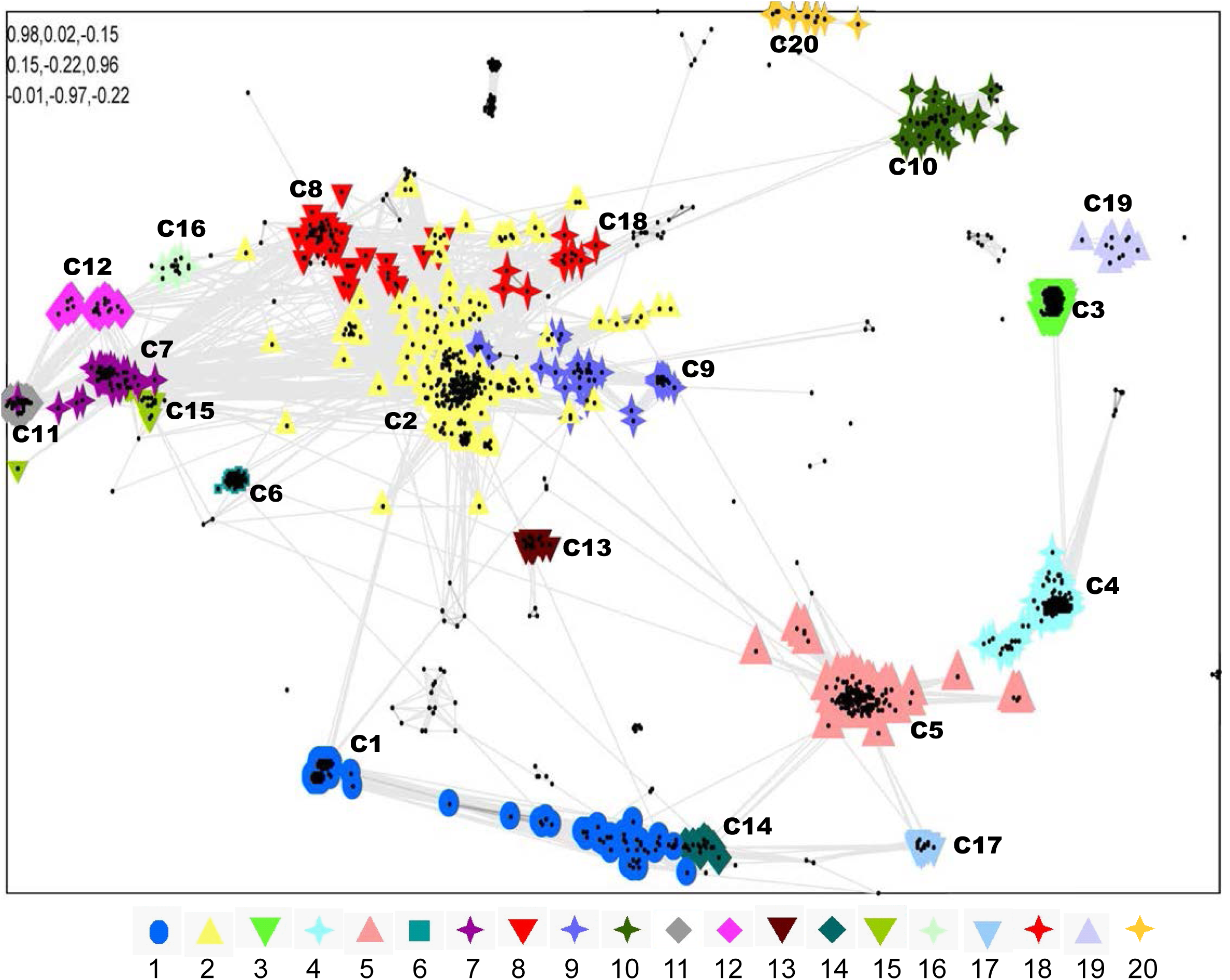
A total of 1823 amino acid sequences of Cap protein of CRESS-DNA viruses were (Supplementary Data 6) used and classified by their pairwise sequence similarity network using CLANS. The clusters were classified using the Network-based method using offset values and global average with a maximum of 10000 in CLANS Toolkit analysis. The *P*-value ≤ 1e^-02^ was used to show the lines connecting the sequences.

**Supplementary Figure 3:**
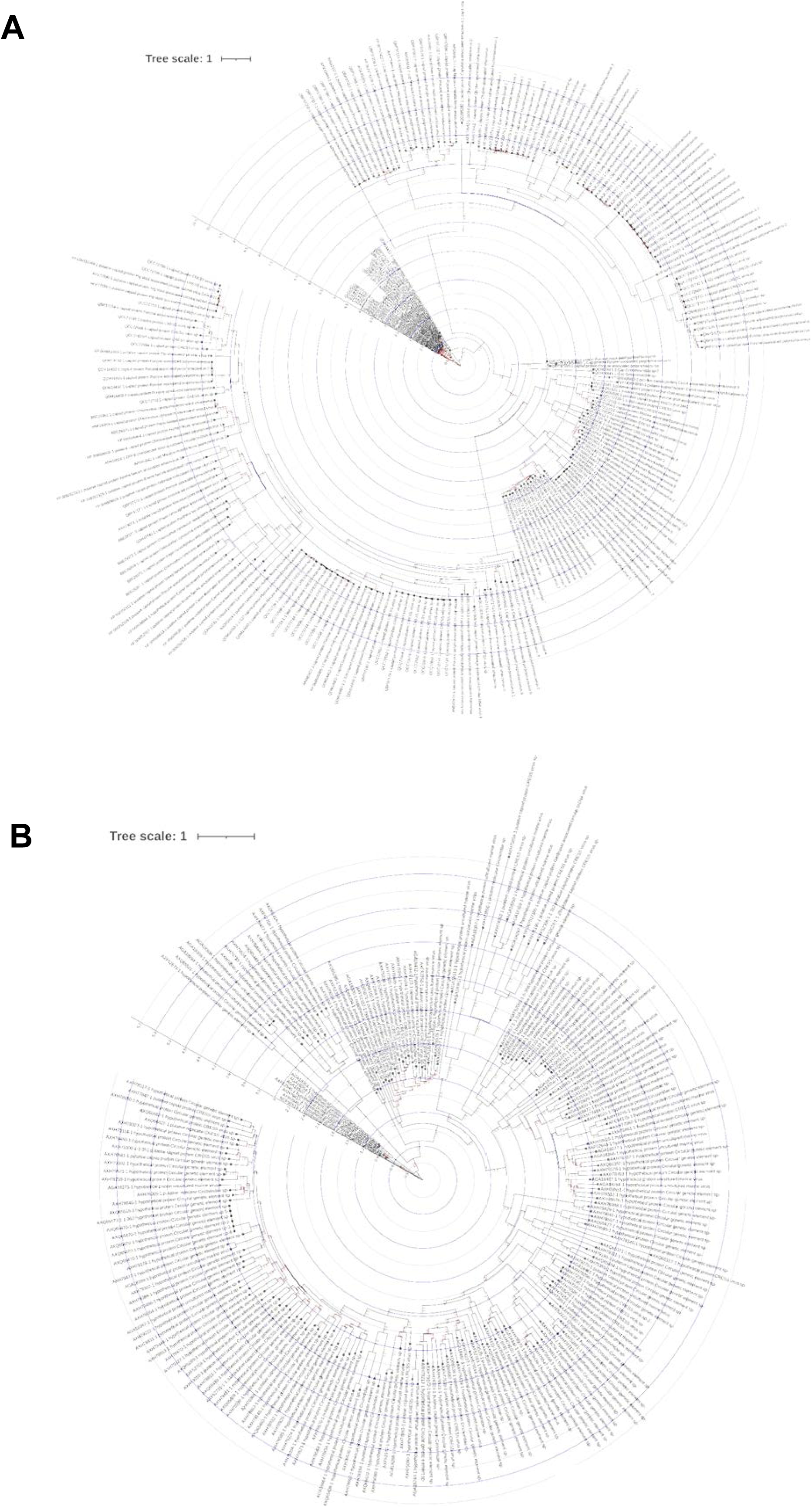
Phylogenetic relationship of CRESS-DNA virus Cap protein cluster C3 and C4. (**A**) Phylogenetic tree depicting the genetic relationship between the CRESS-DNA virus Cap protein cluster C3. The details of sequences in the cluster (**Supplementary Data 7**) and alignment are provided in **Supplementary Data 12**. (**B**) Phylogenetic tree depicting the genetic relationship between the CRESS-DNA virus Cap protein cluster C4. The details of sequences in the cluster (**Supplementary Data 7**) and alignment are provided in **Supplementary Data 13**. The maximum-likelihood method inferred the evolutionary history using the Subtree-Pruning-Regrafting algorithm and bootstrap values in PhyML 3.3_1.

**Supplementary Figure 4:**
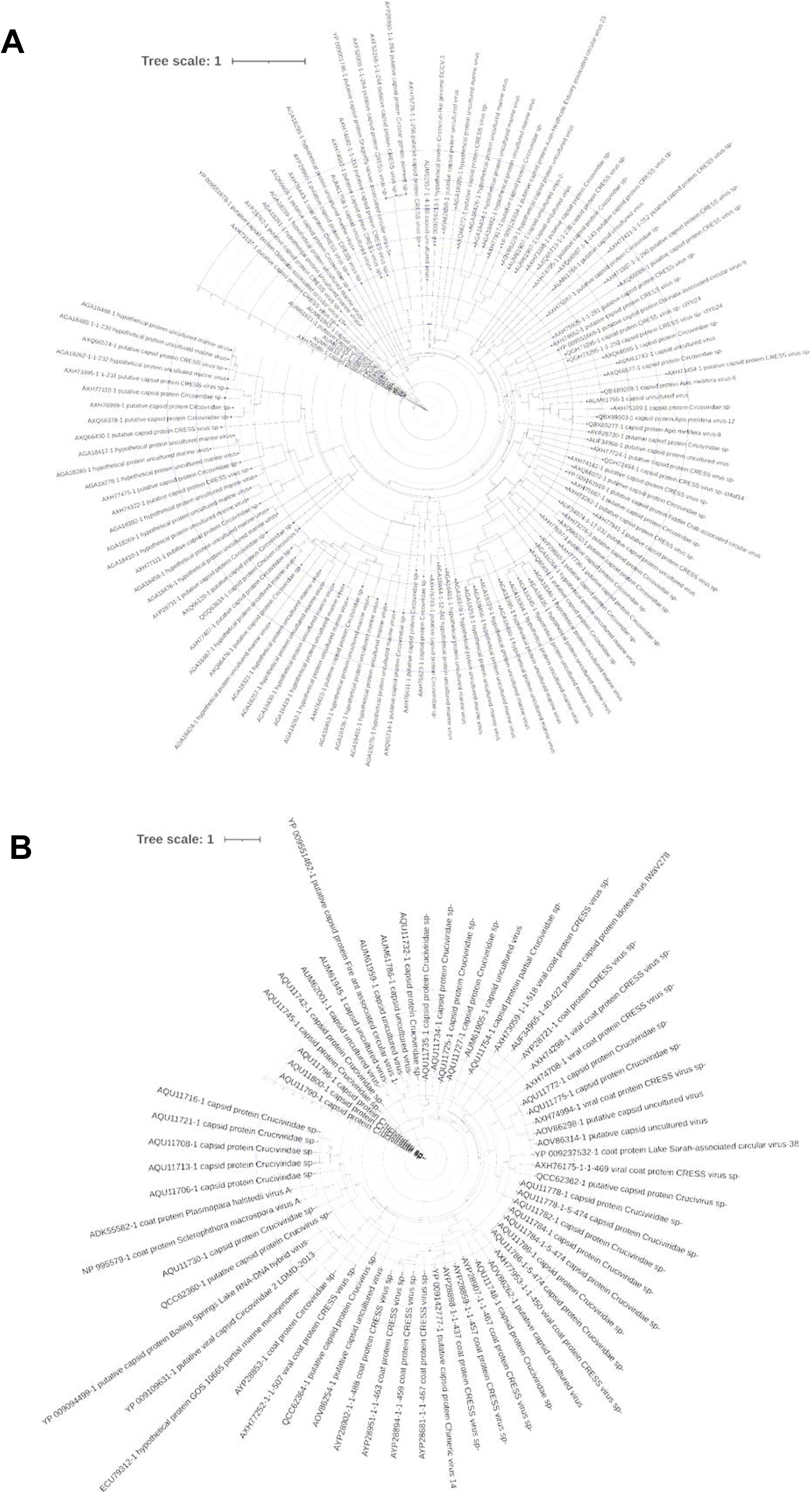
Phylogenetic relationship of CRESS-DNA virus Cap protein cluster C6 and C7. (**A**) Phylogenetic tree depicting the genetic relationship between the CRESS-DNA virus Cap protein cluster C6. The details of sequences in the cluster (**Supplementary Data 7**) and alignment are provided in **Supplementary Data 14**. (**B**) Phylogenetic tree depicting the genetic relationship between the CRESS-DNA virus Cap protein cluster C7. The details of sequences in the cluster (**Supplementary Data 7**) and alignment are provided in **Supplementary Data 15**. The maximum-likelihood method inferred the evolutionary history using the Subtree-Pruning-Regrafting algorithm and bootstrap values in PhyML 3.3_1.

**Supplementary Figure 5:**
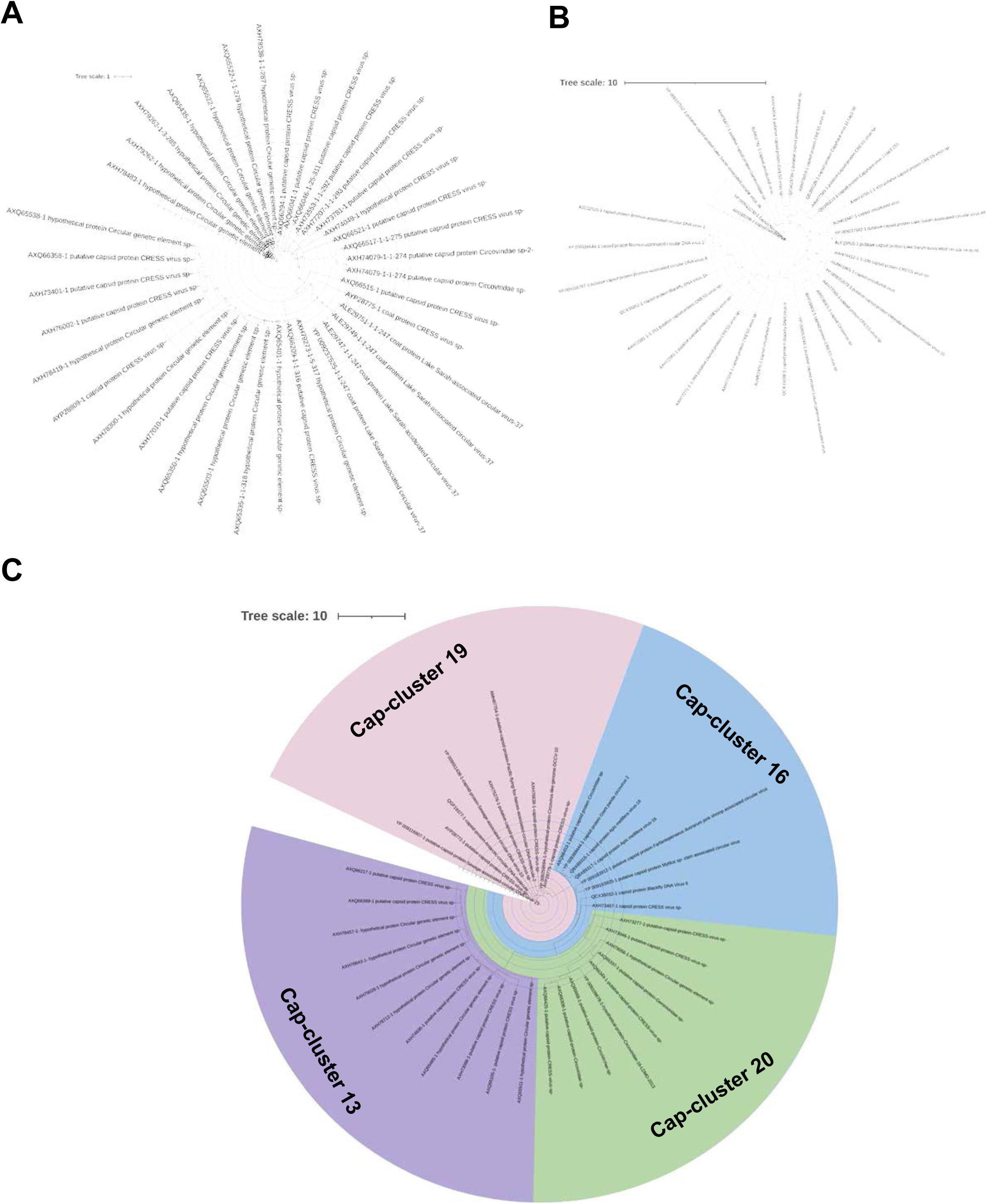
Phylogenetic relationship of CRESS-DNA virus Cap protein cluster C10, C11 C13, C16, C19, and C20. (**A**) Phylogenetic tree depicting the genetic relationship between the CRESS-DNA virus Cap protein cluster C10. The details of sequences in the cluster (**Supplementary Data 7**) and alignment are provided in **Supplementary Data 16**. (**B**) Phylogenetic tree depicting the genetic relationship between the CRESS-DNA virus Cap protein cluster C11. The details of sequences in the cluster (**Supplementary Data 7**) and alignment are provided in **Supplementary Data 17**. (**B**) Phylogenetic tree depicting the genetic relationship between the CRESS-DNA virus Cap protein cluster C13, C16, C19, and C20. The details of sequences in each cluster (**Supplementary Data 7**) and alignment are provided in **Supplementary Data 18**. The maximum-likelihood method inferred the evolutionary history using the Subtree-Pruning-Regrafting algorithm and bootstrap values in PhyML 3.3_1.

## References

1 Kazlauskas, D., Varsani, A., Koonin, E. V. & Krupovic, M. Multiple origins of prokaryotic and eukaryotic single-stranded DNA viruses from bacterial and archaeal plasmids. Nature communications 10, 3425, doi:10.1038/s41467-019-11433-0 (2019).

2 Zhao, L., Rosario, K., Breitbart, M. & Duffy, S. Eukaryotic Circular Rep-Encoding Single-Stranded DNA (CRESS DNA) Viruses: Ubiquitous Viruses With Small Genomes and a Diverse Host Range. Advances in virus research 103, 71–133, doi:10.1016/bs.aivir.2018.10.001 (2019).

3 Krupovic, M. Networks of evolutionary interactions underlying the polyphyletic origin of ssDNA viruses. Current opinion in virology 3, 578–586, doi:10.1016/j.coviro.2013.06.010 (2013).

4 Chow, C. E. & Suttle, C. A. Biogeography of Viruses in the Sea. Annual review of virology 2, 41–66, doi:10.1146/annurev-virology-031413-085540 (2015).

5 Labonte, J. M. & Suttle, C. A. Previously unknown and highly divergent ssDNA viruses populate the oceans. The ISME journal 7, 2169–2177, doi:10.1038/ismej.2013.110 (2013).

6 Ng, T. F. et al. High variety of known and new RNA and DNA viruses of diverse origins in untreated sewage. Journal of virology 86, 12161–12175, doi:10.1128/JVI.00869-12 (2012).

7 Dayaram, A. et al. Diverse circular replication-associated protein encoding viruses circulating in invertebrates within a lake ecosystem. Infection, genetics and evolution : journal of molecular epidemiology and evolutionary genetics in infectious diseases 39, 304–316, doi:10.1016/j.meegid.2016.02.011 (2016).

8 Rosario, K., Schenck, R. O., Harbeitner, R. C., Lawler, S. N. & Breitbart, M. Novel circular single-stranded DNA viruses identified in marine invertebrates reveal high sequence diversity and consistent predicted intrinsic disorder patterns within putative structural proteins. Frontiers in microbiology 6, 696, doi:10.3389/fmicb.2015.00696 (2015).

9 Dayaram, A. et al. Diverse small circular DNA viruses circulating amongst estuarine molluscs. Infection, genetics and evolution : journal of molecular epidemiology and evolutionary genetics in infectious diseases 31, 284–295, doi:10.1016/j.meegid.2015.02.010 (2015).

10 Bistolas, K. S. I., Rudstam, L. G. & Hewson, I. Gene expression of benthic amphipods (genus: Diporeia) in relation to a circular ssDNA virus across two Laurentian Great Lakes. PeerJ 5, e3810, doi:10.7717/peerj.3810 (2017).

11 Blinkova, O. et al. Frequent detection of highly diverse variants of cardiovirus, cosavirus, bocavirus, and circovirus in sewage samples collected in the United States. Journal of clinical microbiology 47, 3507–3513, doi:10.1128/JCM.01062-09 (2009).

12 Krupovic, M. et al. Cressdnaviricota: a Virus Phylum Unifying Seven Families of Rep-Encoding Viruses with Single-Stranded, Circular DNA Genomes. Journal of virology 94, doi:10.1128/JVI.00582-20 (2020).

13 Kazlauskas, D. et al. Evolutionary history of ssDNA bacilladnaviruses features horizontal acquisition of the capsid gene from ssRNA nodaviruses. Virology 504, 114–121, doi:10.1016/j.virol.2017.02.001 (2017).

14 Diemer, G. S. & Stedman, K. M. A novel virus genome discovered in an extreme environment suggests recombination between unrelated groups of RNA and DNA viruses. Biology direct 7, 13, doi:10.1186/1745-6150-7-13 (2012).

15 Roux, S. et al. Chimeric viruses blur the borders between the major groups of eukaryotic single-stranded DNA viruses. Nature communications 4, 2700, doi:10.1038/ncomms3700 (2013).

16 Krupovic, M., Ravantti, J. J. & Bamford, D. H. Geminiviruses: a tale of a plasmid becoming a virus. BMC evolutionary biology 9, 112, doi:10.1186/1471-2148-9-112 (2009).

17 de la Higuera, I. et al. Unveiling Crucivirus Diversity by Mining Metagenomic Data. mBio 11, doi:10.1128/mBio.01410-20 (2020).

18 Yoon, H. S. et al. Single-cell genomics reveals organismal interactions in uncultivated marine protists. Science 332, 714–717, doi:10.1126/science.1203163 (2011).

19 Kazlauskas, D., Varsani, A. & Krupovic, M. Pervasive Chimerism in the Replication-Associated Proteins of Uncultured Single-Stranded DNA Viruses. Viruses 10, doi:10.3390/v10040187 (2018).

20 Simmonds, P. et al. Consensus statement: Virus taxonomy in the age of metagenomics. Nature reviews. Microbiology 15, 161–168, doi:10.1038/nrmicro.2016.177 (2017).

21 Zimmermann, L. et al. A Completely Reimplemented MPI Bioinformatics Toolkit with a New HHpred Server at its Core. J Mol Biol 430, 2237–2243, doi:10.1016/j.jmb.2017.12.007 (2018).

22 Frickey, T. & Lupas, A. CLANS: a Java application for visualizing protein families based on pairwise similarity. Bioinformatics 20, 3702–3704, doi:10.1093/bioinformatics/bth444 (2004).

23 Krupovic, M. & Koonin, E. V. Multiple origins of viral capsid proteins from cellular ancestors. Proc Natl Acad Sci U S A 114, E2401–E2410, doi:10.1073/pnas.1621061114 (2017).

24 Whitley, C. et al. Novel replication-competent circular DNA molecules from healthy cattle serum and milk and multiple sclerosis-affected human brain tissue. Genome announcements 2, doi:10.1128/genomeA.00849-14 (2014).

25 Gorbalenya, A. E., Koonin, E. V. & Wolf, Y. I. A new superfamily of putative NTP-binding domains encoded by genomes of small DNA and RNA viruses. FEBS letters 262, 145–148, doi:10.1016/0014-5793(90)80175-i (1990).

26 Lu, S. et al. CDD/SPARCLE: the conserved domain database in 2020. Nucleic acids research 48, D265–D268, doi:10.1093/nar/gkz991 (2020).

27 Marchler-Bauer, A. et al. CDD/SPARCLE: functional classification of proteins via subfamily domain architectures. Nucleic acids research 45, D200–D203, doi:10.1093/nar/gkw1129 (2017).

28 Marchler-Bauer, A. et al. CDD: NCBI’s conserved domain database. Nucleic acids research 43, D222–226, doi:10.1093/nar/gku1221 (2015).

29 Marchler-Bauer, A. et al. CDD: a Conserved Domain Database for the functional annotation of proteins. Nucleic acids research 39, D225–229, doi:10.1093/nar/gkq1189 (2011).

30 Abrescia, N. G., Bamford, D. H., Grimes, J. M. & Stuart, D. I. Structure unifies the viral universe. Annu Rev Biochem 81, 795–822, doi:10.1146/annurev-biochem-060910-095130 (2012).

31 Greene, L. H. et al. The CATH domain structure database: new protocols and classification levels give a more comprehensive resource for exploring evolution. Nucleic acids research 35, D291–297, doi:10.1093/nar/gkl959 (2007).

32 Krupovic, M. & Bamford, D. H. Double-stranded DNA viruses: 20 families and only five different architectural principles for virion assembly. Current opinion in virology 1, 118–124, doi:10.1016/j.coviro.2011.06.001 (2011).

33 Du, M. Z. et al. The GC Content as a Main Factor Shaping the Amino Acid Usage During Bacterial Evolution Process. Front Microbiol 9, 2948, doi:10.3389/fmicb.2018.02948 (2018).

34 Bohlin, J., Brynildsrud, O., Vesth, T., Skjerve, E. & Ussery, D. W. Amino acid usage is asymmetrically biased in AT-and GC-rich microbial genomes. PLoS One 8, e69878, doi:10.1371/journal.pone.0069878 (2013).

35 Zhao, Y. et al. Analysis of codon usage bias of envelope glycoprotein genes in nuclear polyhedrosis virus (NPV) and its relation to evolution. BMC Genomics 17, 677, doi:10.1186/s12864-016-3021-7 (2016).

36 Wang, L. et al. Genome-wide analysis of codon usage bias in four sequenced cotton species. PLoS One 13, e0194372, doi:10.1371/journal.pone.0194372 (2018).

37 Gun, L., Yumiao, R., Haixian, P. & Liang, Z. Comprehensive Analysis and Comparison on the Codon Usage Pattern of Whole Mycobacterium tuberculosis Coding Genome from Different Area. Biomed Res Int 2018, 3574976, doi:10.1155/2018/3574976 (2018).

38 Tian, H. F. et al. Genetic and codon usage bias analyses of major capsid protein gene in Ranavirus. Infect Genet Evol 84, 104379, doi:10.1016/j.meegid.2020.104379 (2020).

39 Guo, Z., He, Q., Tang, C., Zhang, B. & Yue, H. Identification and genomic characterization of a novel CRESS DNA virus from a calf with severe hemorrhagic enteritis in China. Virus research 255, 141–146, doi:10.1016/j.virusres.2018.07.015 (2018).

40 Moens, M. A. J., Perez-Tris, J., Cortey, M. & Benitez, L. Identification of two novel CRESS DNA viruses associated with an Avipoxvirus lesion of a blue-and-gray Tanager (Thraupis episcopus). Infection, genetics and evolution : journal of molecular epidemiology and evolutionary genetics in infectious diseases 60, 89–96, doi:10.1016/j.meegid.2018.02.015 (2018).

41 Liu, Q. et al. Viral metagenomics revealed diverse CRESS-DNA virus genomes in faeces of forest musk deer. Virology journal 17, 61, doi:10.1186/s12985-020-01332-y (2020).

42 Lemoine, F. et al. Renewing Felsenstein’s phylogenetic bootstrap in the era of big data. Nature 556, 452–456, doi:10.1038/s41586-018-0043-0 (2018).

43 Guindon, S. et al. New algorithms and methods to estimate maximum-likelihood phylogenies: assessing the performance of PhyML 3.0. Systematic biology 59, 307–321, doi:10.1093/sysbio/syq010 (2010).

44 Lemoine, F. et al. NGPhylogeny.fr: new generation phylogenetic services for non-specialists. Nucleic acids research 47, W260–W265, doi:10.1093/nar/gkz303 (2019).

45 Letunic, I. & Bork, P. Interactive Tree Of Life (iTOL) v5: an online tool for phylogenetic tree display and annotation. Nucleic acids research 49, W293–W296, doi:10.1093/nar/gkab301 (2021).

46 Vetrivel, U., Arunkumar, V. & Dorairaj, S. ACUA: a software tool for automated codon usage analysis. Bioinformation 2, 62–63, doi:10.6026/97320630002062 (2007).

47 Puigbo, P., Bravo, I. G. & Garcia-Vallve, S. CAIcal: a combined set of tools to assess codon usage adaptation. Biol Direct 3, 38, doi:10.1186/1745-6150-3-38 (2008).

